# An Eyewear-Based EEG Interface for Temporal-Region Monitoring and Emotion Recognition

**DOI:** 10.64898/2026.11.02.763849

**Authors:** Xiaotao Li, Lining Chai, Shen Xu, Jianguang Ni, Kwok-Fai So

## Abstract

Conventional electroencephalography (EEG) devices are often impractical for ambulatory settings due to their reliance on specialized hardware and expert operation. To address this, we developed a user-friendly eyewear-based EEG system featuring dry titanium electrodes positioned approximately 5 mm inferoposterior to the T7/T8 sites (near the ears), with metal nose pads serving as references. We evaluated the system across 103 recordings spanning five paradigms: alpha reactivity, blink-related transients, fatigue, attention, and video-elicited emotion. Following quality control, 93.2–100.0% of epochs were retained. The system demonstrated high sensitivity to ocular transients, showing significant amplitude increases during the blinking task. The emotion task yielded the most robust state-related features, with statistically significant differences in differential entropy across negative, neutral, and positive video categories, achieving balanced accuracies of 0.882 for two-class and 0.667 for three-class prediction. Despite challenges at the multiple testing correction stage, the alpha reactivity and fatigue contrast tasks were still noteworthy to some extent. The attention paradigm revealed a consistent increase in differential entropy during focused states, and the eyewear system outperformed the Muse headband in group-held-out classification. These findings provide strong evidence for the feasibility of temporal-region eyewear EEG acquisition, highlighting its potential applications in the real-world, particularly for emotion recognition.

## I. Introduction

ELECTROENCEPHALOGRAPHY(EEG) provides a physiological window into brain activity and has been widely used in sleep-phase classification [1], epilepsy detection [2], and cognitive assessment [3], [4]. Due to its inherent safety, recently, EEG technology has witnessed a marked surge in popularity as a tool of non-invasive brain-computer interface (BCI) that facilitates the translation of brain activity into a format that can be interpreted by external devices [5], [6]. The decoding methodologies employed by this technology have also undergone significant advances, largely attributable to the integration with deep neural networks [7], [8]. A classical EEG cap, one of the most widely used brainwave-recording devices, is characterized by multiple channel sizes; however, its daily applications have been limited due to complex procedures, time-consuming preparation, and strict operating conditions of traditional recording systems [6], [9]. Some reliance on conductive gel has been observed to result in user discomfort and the necessity of training for its proper operation [4]. Thus, data acquisition is often confined to controlled experimental conditions, which limits its application for long-term ambulatory recording [9]. This limitation is particularly salient in some scenarios, such as fatigue-related monitoring and emotional valence tracking in vehicles [10], where portability, comfort, and robustness to real-world conditions are usually essential [6]. Consequently, there is an emerging demand and concomitant development trend for a wearable, user-friendly, and operationally accessible EEG device, particularly within the domain of non-invasive BCI [5].

In the context of ongoing research endeavors aimed at the development of portable EEG devices, notable progress has been observed in the field [11], [12]. For instance, studies have been conducted on portable BCI devices like the Muse headband [13], cEEGrid [14], and Emotiv EPOC [15]. These studies have yielded advances in the realm of non-invasive BCI technology and the evaluation of mental health [16], [17]. However, most existing devices still suffer from some typical shortcomings, including (1) excessive weight, insufficient comfort, and unappealing aesthetics resulting in low user acceptance [6]; (2) devices’ reliance on external power sources, making long-term continuous collection impossible [18]; (3) most are confined to simple entertainment scenarios and rarely employed in advanced uses such as diagnostic and therapeutic yield [19]. Although a few studies have attempted EEG devices’ use for medical treatment [20], neurofeedback therapy [21], and even home-based monitoring [16], there is a preliminary promise that no mature device solutions or validation systems have been established, leaving significant room for further optimization and improvement [5].

The development of a lightweight, wearable, and user-friendly EEG device with stable recording capabilities, along with a rigorous evaluation of its signal sensitivity across controlled paradigms, are crucial steps towards the practical implementation of non-invasive BCI in the real world. In recent years, ear-based EEG recording has garnered attention for its potential in long-term recording and the rapid development of portable BCI devices [22], [18]. Among these, the cEEGrid, a representative periauricular flexible electrode array [23], captures EEG mainly from the temporal lobe projection area through multi-channel sensor units around the auricle [24], showing some promise for sleep monitoring and epilepsy detection in naturalistic settings [26]. In the 10-10 electrode placement system, T7 and T8 are located over the bilateral temporal lobes, namely T3 and T4 in the 10-20 system, which are the closest sites to our ears, overlied some parts of middle and inferior temporal gyri regions involved in multisensory integration and emotional processing [24], [27]. Several studies have reported that EEG signals especially collected at T7/T8 sites provided useful information mainly from the temporal lobes and show consistency with homologous temporal electrodes of traditional EEG caps [25], [26], [28]. In this study, we propose lightweight, eyewear-based EEG system to capture EEG signals inferoposterior to the T7/T8 sites. This device integrates EEG electrodes into the glasses frame using dry titanium electrodes [6], [29], with the temple-arm contacts positioned to ears inferoposterior to the T7/T8 sites and metal nose pads serving as the EEG references [30], [31]. The device, which bears a resemblance to conventional eyewear, is designed to facilitate unobtrusive and operator-light EEG recording in routine daily activities [32]. However, the efficacy of this eyewear-based EEG system for extended real-world monitoring remains to be substantiated through further validation.

In order to evaluate our system as an EEG acquisition platform, five established experimental paradigms were utilized. These paradigms included (1) eyes-open versus eyes-closed alpha reactivity, (2) normal resting versus voluntary blinking, (3) awake versus fatigued resting state, (4) distracted versus focused attention, and (5) negative, neutral, and positive emotion elicitation [26], [28]. Available de-identified recordings were organized into recording units within each paradigm. Raw EEG signals were preprocessed and summarized using time-domain, spectral, and entropy-based features [33]; pre-specified primary contrasts were tested at the recording-unit level using paired sign-flip permutation tests with false-discovery-rate correction; and group-held-out classification was used as a predictive sensitivity analysis within each paradigm [34]. We further used wavelet-resolved analysis to examine frequency-specific effects and benchmarked sensitivity against that of Muse headband recording and the dataset collected by an EEG cap with 62 channels.

## II. Data Acquisition And Experiments

### A. Experimental Equipment and Related Materials

Fig. 1 provides a synopsis of the hardware and acquisition pathway. The dry titanium electrodes positioned at the temple tips made contact with the skin approximately 5 millimeters inferoposterior to T7 and T8 points, with the two titanium nose pads serving as the reference points [30], [31]. The analog front end exhibited an input impedance of 47 MΩ, 16-bit analog-to-digital conversion capability, and a device-reported signal-to-noise ratio of 74.5 dB. EEG signals were sampled at 512 Hz, packetized by the microcontroller, and transmitted through Bluetooth Low Energy (BLE 4.0/5.0) to acquisition software running on a laptop computer. The device specification indicated a wireless latency of no more than 50 milliseconds at 512 Hz.

**FIG. 1.**
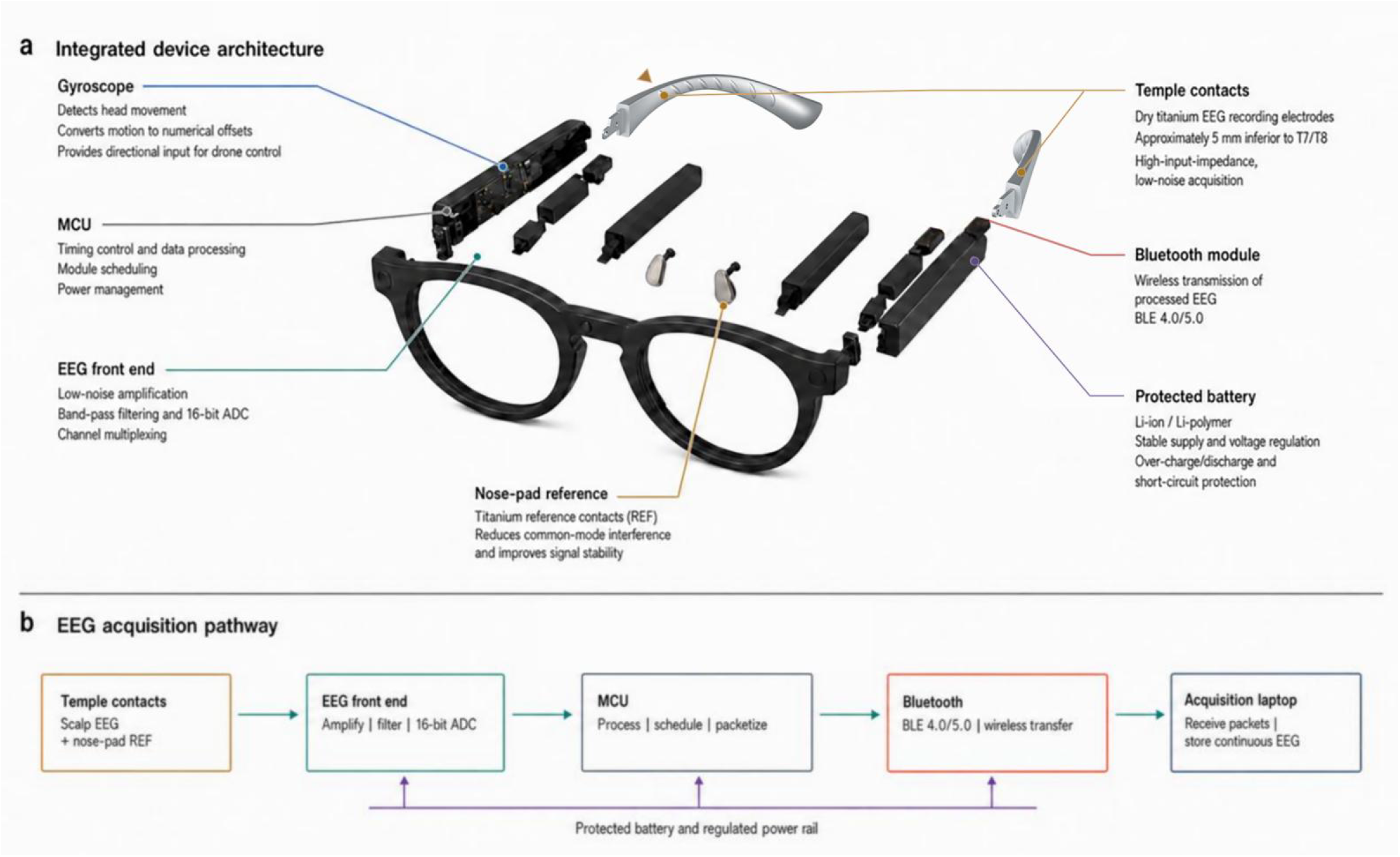
Eyewear-based EEG System architecture, Acquisition pathway. (a) Exploded view of the glasses frame-integrated electronics, including dry titanium temple contacts positioned approximately 5 mm inferoposterior to T7/T8 sites, titanium nose-pad references, the EEG front end, microcontroller, protected battery, bluetooth module, and gyroscope. (b) Acquisition pathway from scalp-potential sensing to amplification, digitization, packetization, BLE transfer, and storage on the acquisition laptop; the integrated motion channel was not present in the archived EEG files and was not analysed yet.

The study utilized the eyewear-based EEG system, together with one laptop running the acquisition software, and another laptop presenting some visual stimuli and other tasks, including a neutral Windows desktop image, an 8 × 8 Schulte grid, emotion-eliciting clips from the SEED dataset [28], and also a questionnaire of the Karolinska Sleepiness Scale (KSS) [35]. A modified BCI-controlled drone was utilized separately to test the feasibility of the eyewear EEG glasses frame to run a drone with the user’s blinking and focused action code.

Following the completion of the aforementioned preparatory procedures, the acquisition laptop successfully received BLE packets; it should be noted that the utilization of a smartphone was not a requisite component of this experimental setup. Each continuous recording was stored as a UTF-8 text file containing one numeric sample list. The initiation of the file was designated as 0 seconds (s), as the archive failed to incorporate synchronized event markers or absolute timestamps. A secondary laptop furnished the screen-based paradigms illustrated in Fig. 2.

**FIG. 2.**
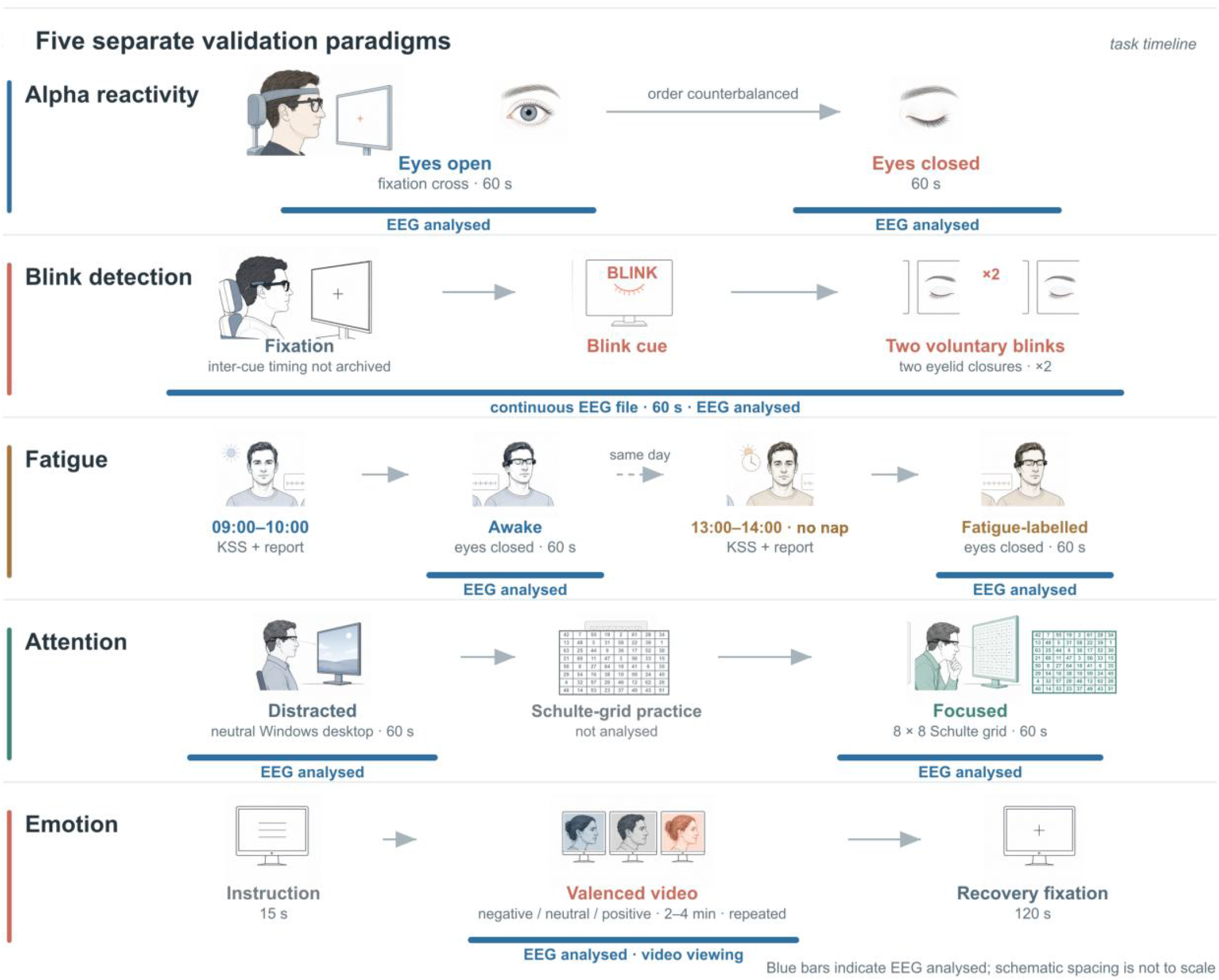
Experimental paradigms. Five separate validation paradigms showing: (a) Alpha reactivity, (b) Blink detection, (c) Fatigue, (d) Attention, (e) Emotion timelines. Blue bars indicate the continuous EEG files included in analysis; schematic spacing is not to scale.

Prior to the initiation of the recording, the temporal contact sites were meticulously cleansed using a combination of wet and dry tissues. The subject’s hair was carefully tidied to ensure it did not come into contact with the titanium electrodes, especially for females with long hair. The glasses frame was also adjusted to ensure optimal contact without exerting additional pressure. The hardware contains a six-axis motion sensor to detect the motion activities from the head, eyes, and so on. The exported EEG values also lacked a calibration factor for conversion to microvolts.

Consequently, absolute-amplitude panels report device output units (a.u.) instead of physical voltage. The analysis archive was composed of five task folders: alpha reactivity, blink detection, fatigue, attention, and emotion recognition.

The labels of files and folders were parsed into the following categories: task, condition, participant label, device-version label (where present), and source file. The definition of a recording unit was established as the smallest within-paradigm unit that could be paired across conditions. All inferences were conducted within the paradigm. The aggregation of summaries was constrained to file counts, recording-unit counts, and quality-control retention. This design precluded the occurrence of substantial blink transients or extended emotion recordings from exerting an influence on cross-paradigm feature or classification claims.

### B. Experimental Design

Five validation paradigms were administered as separate experiments (Fig. 2). The analyzed alpha, blink, fatigue, and attention files each contained approximately 60 s per condition; the emotion files contained 111.4 to 120.0 s of continuous clip viewing. Each paradigm was designed to assess a specific acquisition target, including alpha reactivity, blink detection, fatigue-related state, attention engagement, and video-elicited emotion classification.

The five experimental paradigms were retained as separate validation experiments. This distinction was pivotal in the revised analysis, as the paradigms did not assess the same physiological construct. The experimental design includes a series of tests designed to elicit different forms of ocular and behavioral responses. These include a comparison of eyes-open versus eyes-closed recording tests, which measure spectral alpha reactivity. Additionally, there is a comparison of normal resting versus voluntary blinking tests, which measure large ocular transients. The study also includes a comparison of awake versus fatigued resting recording tests, which measure a fatigue state shift. Furthermore, there is a comparison of distracted versus focused attention tests, which measure attention engagement. Finally, the design includes a series of tests that elicit emotional states through negative, neutral, and positive video elicitation.

For each paradigm, at least one contrast and one primary feature were defined for inference. Alpha oscillation utilized the eyes-closed minus eyes-open contrast (Fig. 2a); blinking employed voluntary blinking minus normal resting (Fig. 2b); fatigue employed fatigued minus awake (Fig. 2c); attention employed focused minus distracted (Fig. 2d); while emotion employed positive minus negative as the primary binary contrast, with an additional three-condition negative-neutral-positive test (Fig. 2e). This prevented exploratory features from being presented as if they had been selected prior to examining the data.

#### 1) Alpha Oscillation with Closed Eyes

The alpha-reactivity experiment consisted of two segments: the first segment was 60 s of relaxed eyes-open viewing, and the second segment was 60 s with eyes closed. The condition order was reversed across acquisition blocks to reduce order dependence. The final archive contained eight matched recording units, with the eyes-open/eyes-closed pairs in each unit.

The pre-specified endpoint of interest was the recording-unit median alpha/beta ratio [36]. The selection of this ratio was predicated on two factors: first, the anticipated increase in alpha activity during periods of eye closure [37], and second, the reduced sensitivity of within-unit ratios to variations in between-session amplitude scaling [38].

#### 2) Blinking Detection

The blink experiment was designed to compare normal rest with prompted blinking [39]. During periods of rest, participants were instructed to remain motionless and to minimize blinking. During the prompted condition, the word “BLINK” appeared on the stimulus laptop, and participants were instructed to produce two consecutive blinks per cue. Despite the incorporation of extended task periods and intermissions as part of the acquisition protocol, the archived files utilized in this study contained 60 s per condition.

The blink transients were retained due to their status as the target signal, as opposed to their treatment as artifacts. The pre-specified endpoint of interest was the median peak-to-peak amplitude of the recording unit [40], [41].

#### 3) Fatigue Contrast

The prevalence of fatigue was assessed in participants who habitually took a midday nap. The awake recording was obtained between 9:00 am and 10:00 am, whereas the fatigue-labelled recording was obtained between 13:00 pm and 14:00 pm without a nap. During the acquisition process, the KSS ratings and self-report were utilized to verify their states [35]. Participants were instructed to rest with their eyes closed during each 60-s recording that was analyzed.

The pre-specified endpoint was differential entropy [35], [42]. Individual KSS values, psychomotor-vigilance measures, and circadian covariates were included in the analysis archive [35]. However, questionnaire-feature correlations were not described here, and the fatigue contrast is reported as exploratory rather than diagnostic.

#### 4) Attention Engagement

Compared with looking around a neutral desktop image as control group, the attention experiment was conducted using an 8 × 8 Schulte-grid task. Participants initially engaged in a preparatory phase, during which they familiarized themselves with the Schulte-grid task. Thereafter, they proceeded to the execution of the focused condition, while EEGs were concurrently recorded. The analyzed archive contained one 60-s file per condition for each matched unit [43].

The pre-specified endpoint was differential entropy [3], [33]. The classification process was constrained to the attention paradigm. A Muse headband dataset that was acquired separately with the Schulte-grid task was analyzed with the same feature family and group-held-out validation to provide a device control [13], [44].

#### 5) Emotion Recognition

The elicitation of emotion was achieved through the utilization of negative, neutral, and positive clips derived from the SEED dataset [28], [45]. Each clip was preceded by a 15-s instruction period and followed by 120 s of recovery. EEG signals were recorded continuously during the 2-4 minute clips. The archived files available for analysis contained 111.4-120.0 seconds per condition. Participants were instructed to minimize movement and refrain from speaking during the recording session.

The pre-specified binary contrast was defined as positive minus negative differential entropy [45], [46]. A three-condition Friedman test was utilized to assess modulation across negative, neutral, and positive recordings [28].

### C. Participants

Informed consent was obtained from all individual participants included in the study. The analysis employed de-identified participant or recording labels from the folder structure. A unit entered a paired contrast only when both conditions were available; emotion units entered the three-condition analysis only when negative, neutral, and positive files could be matched.

The original cohort comprised fifteen participants (20-47 years; 6 females and 9 males). Due to the unavailability of age and gender data for the additional files, demographic summaries are based on the original cohort. Each analysis reports its task-specific number of matched recording units.

## III. Method

### A. Pre-processing and Feature Extraction

All participants have gotten the informed consent before they performed the tasks of five paradigms, and all data were de-identified prior to analysis.. Each text file was parsed into a numeric time series and checked for missing or non-finite samples. The signals were subjected to a linear detrending procedure, followed by a notch filter at 50 Hz (Q = 30). Subsequently, a band-pass filter was employed, spanning from 0.5 to 60 Hz, with a fourth-order zero-phase Butterworth filter. The recordings were segmented into 2-s epochs, with each epoch separated by a 1-s interval [47].

Zero-phase filtering was found to be an effective method for preserving temporal alignment while suppressing baseline drift and line noise. The archive contained one EEG series per file, precluding the application of independent component analysis, spatial filtering, cross-channel re-referencing, and T7-T8 coherence.

The epoch quality was evaluated within the recording using peak-to-peak amplitude [2], root-mean-square amplitude, and line length. Epochs that demonstrated an absolute robust z-score greater than 8 on any metric were considered non-significant and were, consequently, rejected. In instances where this rule identified more than 20% of a recording, rejection was limited to the highest 20% of the combined robust-artifact score. The retention of blinks occurred due to the utilization of ocular transients as the target signal.

The time-domain features encompassed variance, mean energy, root-mean-square amplitude, peak-to-peak amplitude, line length, zero-crossing rate, and Hjorth activity, mobility, and complexity [48], [49]. Welch spectra yielded relative delta, theta, alpha, beta, and gamma power, alpha/beta and theta/beta ratios, spectral entropy, and dominant frequency [50]. The calculation of differential entropy was derived from epoch variance [45], [46].

The features were calculated for each epoch and subsequently summarized by the median within each recording unit and condition. This hierarchy was designed to retain temporal detail for visualization purposes while ensuring that overlapping epochs were not erroneously treated as independent participants in the analysis. Power spectra and zero-time trajectories were descriptive and did not define additional confirmatory hypotheses.

For each paradigm, one endpoint was pre-specified: the alpha/beta ratio for alpha reactivity, the peak-to-peak amplitude for blinking, and the differential entropy for fatigue, attention, and emotion. The remaining features were subjected to sensitivity analyses.

### B. Morlet Wavelet Analysis

The exploratory time-frequency analysis employed the accepted non-overlapping 2-s epochs. Subsequent to the preprocessing stage, the signals were resampled at a rate of 128 Hz and decomposed with complex Morlet wavelets ranging from 2 to 45 Hz in increments of 1 Hz. The number of cycles was determined by dividing the frequency by four, with a range constrained to 2-8 cycles. Power was then averaged over the central 0.25-1.75 s of each epoch, normalized to total 2–45-Hz power, and log10 transformed [49].

The relative wavelet power was summarized for delta (2-4 Hz), theta (4-8 Hz), alpha (8-13 Hz), beta (13-30 Hz), and gamma (30-45 Hz) using non-overlapping frequency-bin boundaries [50]. The effects of paired conditions were examined at the level of the recording unit, with FDR correction applied across the 25 task-by-band tests. The absence of KSS correlation and bilateral coherence analysis is attributable to the unavailability of participant-level KSS values and synchronized bilateral channels.

### C. Statistical Analysis and Classification

The recording unit, rather than the epoch, was the primary inferential unit. The condition medians were then paired within each unit and contrasted separately for each paradigm.

Median aggregation has been demonstrated to effectively mitigate the impact of brief residual transients and session-specific amplitude excursions. Epoch distributions remained discernible in the trajectory and quality-control panels.

Two-sided sign-flip permutation tests were used to assess paired contrasts, with exhaustive enumeration for up to 12 units and 100,000 random sign patterns otherwise. A plus-one correction was applied to the permutation p values. The mean paired differences were accompanied by 10,000-resample bootstrap 95% confidence intervals and Cohen’s d. The Benjamini-Hochberg false-discovery-rate correction was implemented within each paradigm-specific feature family. The three-condition emotion analysis employed a Friedman test.

The classification process was conducted within the paradigm, employing leave-one-participant-out or leave-one-recording-unit-out validation, as dictated by the available labels. All epochs from the held-out unit were excluded from the training process. The efficacy of standard regularized logistic regression and random forests was appraised at the subject-condition level. The primary objective was to achieve balanced accuracy, with secondary objectives including binary F1 and AUC, as well as three-class macro-F1 and one-versus-one macro AUC.

The implementation of group-held-out validation effectively prevented adjacent windows from the same recording from appearing in both the training and test sets. The classification process was approached as a predictive sensitivity analysis, with the understanding that it did not supplant the established methodology of recording-unit paired inference.

For the Muse benchmark, focused and distracted recordings were processed at 256 Hz with the same filtering family, 2-s epochs, 1-s overlap, feature set, and leave-one-subject-out random-forest analysis [44]. The TP09/TP10 channel was identified as the primary channel due to its proximity to the temporal contact point of the EEG glasses frame. Within-device standardized effects and held-out metrics were used for the purpose of comparison because absolute entropy is contingent upon device scaling [27].

The SEED dataset, captured with a 62-channel ESI NeuroScan System, was re-analyzed using only T7 and T8. Sessions, films, and sliding windows were aggregated within participants before inference. Differential entropy was the primary feature; T7 and T8 were also analyzed separately as sensitivity checks [28],. Paired contrasts used exact sign-flip permutation tests and 10,000-resample bootstrap 95% CIs. Classification used random forests with leave-one-participant-out validation; window-level features were first summarized to the trial level and then to the participant-condition level [49].

The analyses were implemented in Python 3.10, using the NumPy 1.26.4, SciPy 1.15.3, pandas 2.3.3, scikit-learn 1.7.2, MNE-Python 1.6.1, and Matplotlib 3.10.9 libraries. The random seed was fixed at 20260622 for the primary analysis and 20260713 for the wavelet analysis.

## IV. Results

A re-analysis of the eyewear-based EEG system was conducted using 103 single-channel recordings across five paradigms (Fig. 1 and 2). The alpha oscillation subset contained 16 files from eight recording units; blinking, fatigue and attention detection respectively contained 12 files from six recording units, and emotion classification contained 51 files from 17 recording units (see Fig. S1). The majority of the recordings demonstrated continued usability following robust epoch rejection, in which the mean retention was 93.2% for alpha reactivity, 100.0% for blinking, 99.7% for fatigue, 96.6% for attention, and 96.7% for emotion (Fig. S1-C). Representative traces across five paradigms are shown in the device output units recorded from our eyewear-based EEG system since the archive did not contain a microvolt calibration factor (Fig. 3) [51].

**FIG. 3.**
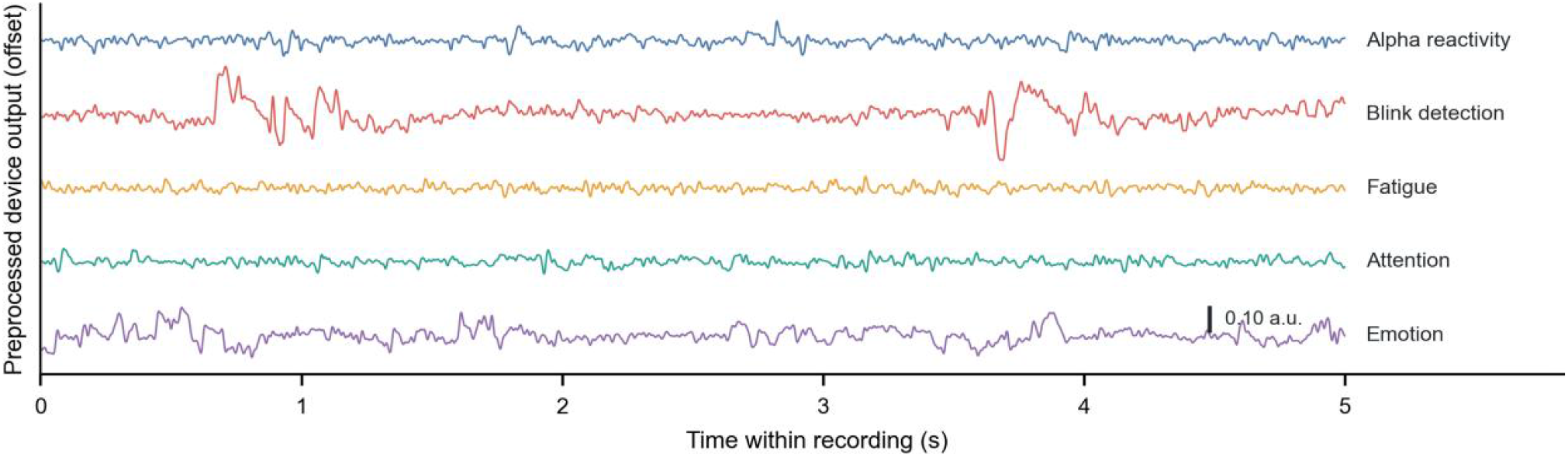
Representative 5-s pre-processed traces in device output units recorded from our eyewear-based EEG system. The vertical bar denotes 0.10 a.u. The archive did not provide a conversion factor to microvolts.

### A. Alpha Oscillations with Eyes Closed

The eyes-open and eyes-closed files were analyzed using the alpha oscillation paradigm [36], [37]. An increase in the alpha/beta ratio was observed during periods of eye closure in six of the eight paired recording units (Fig. 4a). The mean paired difference was 0.3893, with a 95% bootstrap confidence interval ranging from 0.0962 to 0.7152 (Table S1). The paired effect remained in the expected direction but exhibited a little heterogeneity (n = 8, Cohen’s dz = 0.80; FDR q = 0.2588, Fig. 3a and Table S1). These findings are consistent with the spectral profile within the paradigm (Fig. 4b). The eyes-closed condition also exhibited a higher alpha/beta trajectory on average (Fig. 4c). The within-paradigm classifier achieved a subject-condition balanced accuracy of 0.625 and an AUC of 0.766 for the separation of eyes-closed from eyes-open recordings (Fig. 4d and Table 1). Hence, the initial result is understood as a positive-control physiological check [44], rather than as a state classifier that has been fully optimized. This provides support for the validity of our eyewear system as an EEG acquisition system.

**FIG. 4.**
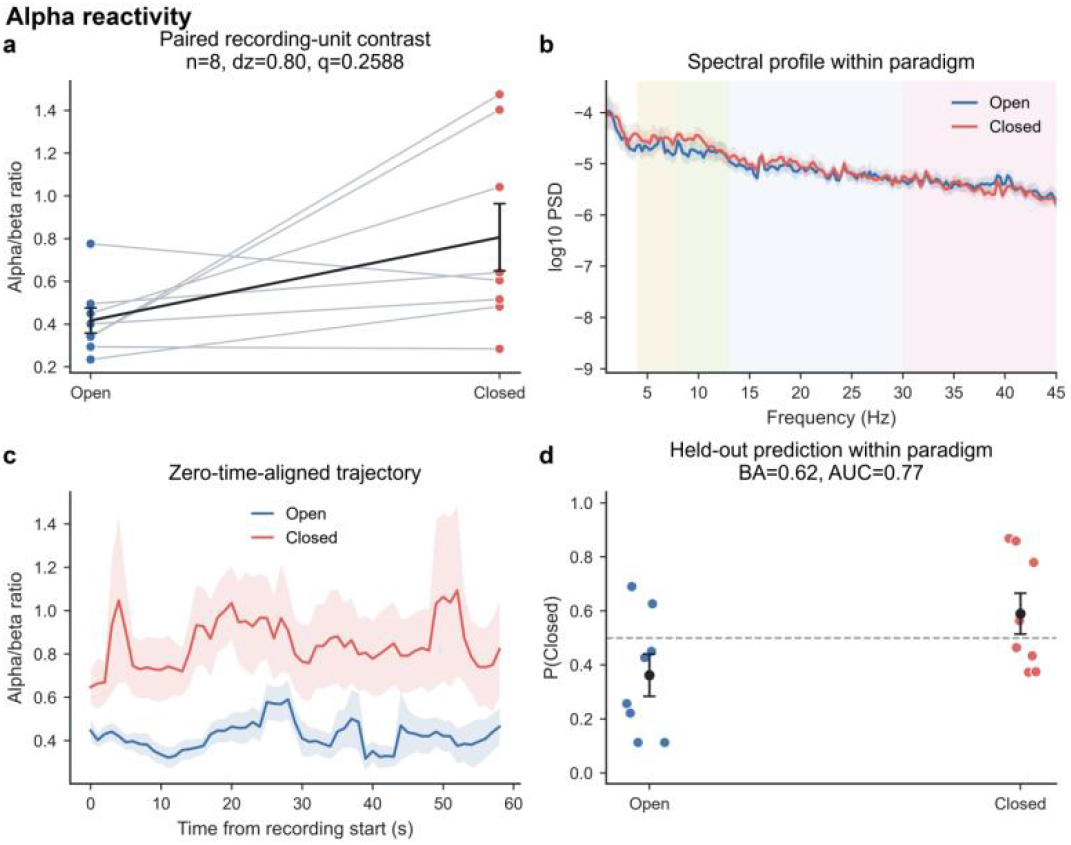
Alpha reactivity. (a) Paired recording-unit alpha/beta ratios (n = 8; closed-open mean difference, 0.3893; 95% CI, 0.0962 to 0.7152; dz = 0.80; sign-flip p = 0.0661; FDR q = 0.2588). (b) Within-paradigm power spectra. (c) Zero-time-aligned alpha/beta trajectories. (d) Group-held-out predictions (balanced accuracy, 0.625; AUC, 0.766). Lines in (a) connect matched units; black symbols show group means and bootstrap 95% confidence intervals.

**TABLE I.** Within-Paradigm Random Forest Performance Under Leave-One-Participant-Out Validation.

| Paradigm | Prediction task | Groups | Test units | Balanced accuracy | F1/macro-F1 | AUC |
|---|---|---|---|---|---|---|
| Alpha reactivity | Closed vs Open | 8 | 16 | 0.625 | 0.571 | 0.766 |
| Blink detection | Blink vs Normal | 6 | 12 | 0.917 | 0.909 | 0.972 |
| Fatigue | Fatigued vs Awake | 6 | 12 | 0.750 | 0.727 | 0.722 |
| Attention | Focused vs Distracted | 6 | 12 | 0.833 | 0.833 | 0.944 |
| Emotion 2 classes | Positive vs Negative | 17 | 34 | 0.882 | 0.889 | 0.976 |
| Emotion, 3 classes | Negative vs Neutral vs Positive | 17 | 51 | 0.667 | 0.669 | 0.815 |

### B. Blinking Detection

The blinking phenomenon elicited one of the most evident positive-control effects. Peak-to-peak amplitude exhibited an increase in all six paired units (mean difference, 0.3404 a.u.; 95% CI, 0.1983 to 0.4741; dz = 1.76; sign-flip p = 0.0462; FDR q = 0.0497, Fig. 5a and Table S1). The random forest classifier achieved a subject-condition balanced accuracy of 0.917 and an AUC of 0.972 for the classification of blink versus normal-rest recordings (Fig. 5d and Table 1). The result is of practical importance for artifact annotation and device interaction [52], though this phenomenon does not fit neatly into the paradigm of a cortical brain state, as the dominant signal source is the blink-related transient itself [39]. Thus, reporting blinking on its own preserves its value as a hardware sensitivity test without overstating the evidence for cognitive-state decoding [39], [40]. As shown in Video S2, in addition, there was a remarkable BCI function to control the roll action of a drone ball [40], which resulted from intentional blinking captured by our EEG system.

**FIG. 5.**
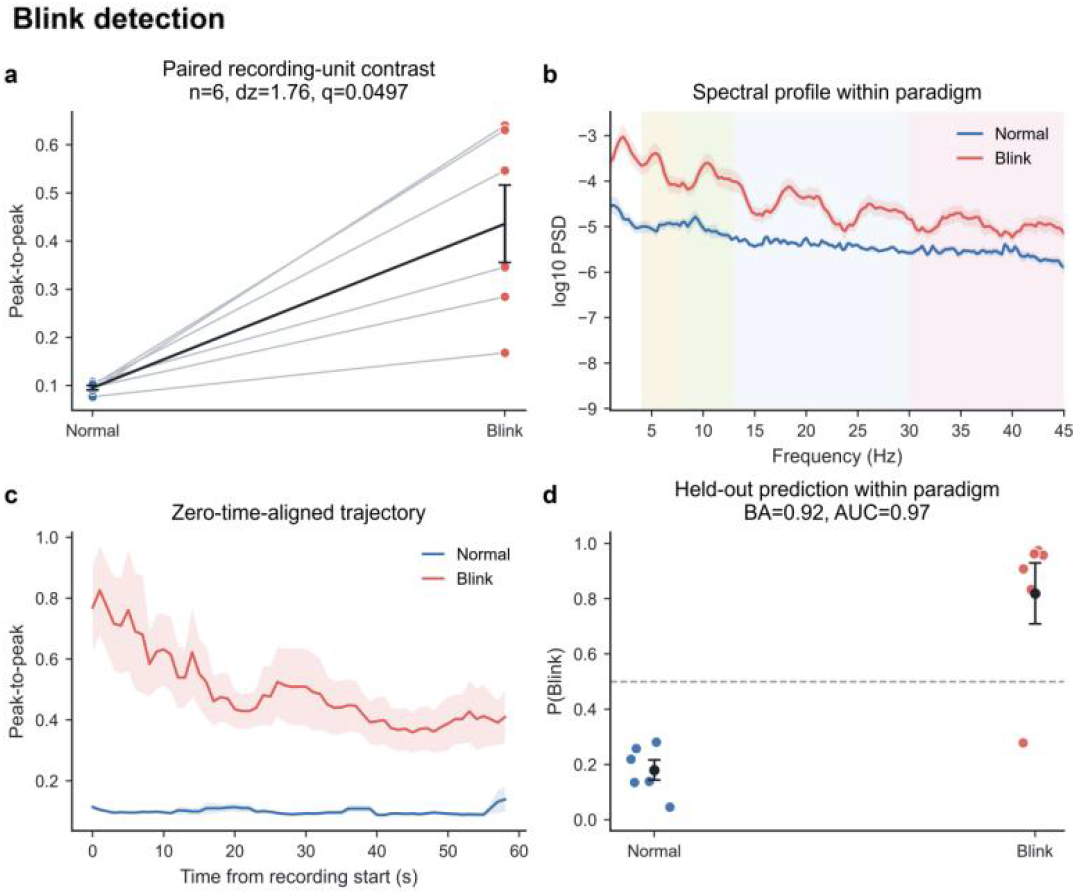
Blink detection. (a) Paired peak-to-peak amplitude (n = 6; blink – normal mean difference, 0.3404 a.u.; 95% CI,0.1983 to 0.4741; dz = 1.76; sign-flip p = 0.0462; FDR q = 0.0497). (b) Within-paradigm power spectra. (c) Zero-time-aligned trajectories. (d) Group-held-out predictions (balanced accuracy, 0.917; AUC, 0.972).

### C. Fatigue Detection

A comparative analysis of the mean paired difference revealed that the mean value for differential entropy in the fatigued recordings was lower than that observed in the awake recordings [35], with a mean difference of −0.1478 (Table S-1). The 95% confidence interval for this mean difference was calculated to be from −0.3462 to −0.0158 (Table S-1). The data were also found to be consistent with a negative slope (dz = −0.63) (Fig. 6a). The sign-flip p-value was 0.1077 (Table S-1), and the corrected q-value was 0.1256 (Fig. 6a). Fatigued recordings also showed exploratory increases in theta and alpha power, while a decrease happened in high-frequency power, especially at gamma power (dz = +0.94, +1.00, and −1.26 for theta, alpha, and gamma, respectively; FDR q = 0.078, Fig. 6b and Fig. 9). The fatigue-related trajectory was consistent with fatigue being slower and more dimensional compared with that of the awake state (Fig. 6c), but it was less sharply separated than the blink or emotion trajectories. The subject-condition random forest classifier achieved a balanced accuracy of 0.750 and AUC of 0.722 (Fig. 6d and Table 1). This performance indicates modest within-dataset separability, but it is not sufficient for a strong claim of fatigue monitoring. Thus, a more rigorous fatigue study would require larger paired samples, longer recordings, or higher performance-based fatigue labels.

**FIG. 6.**
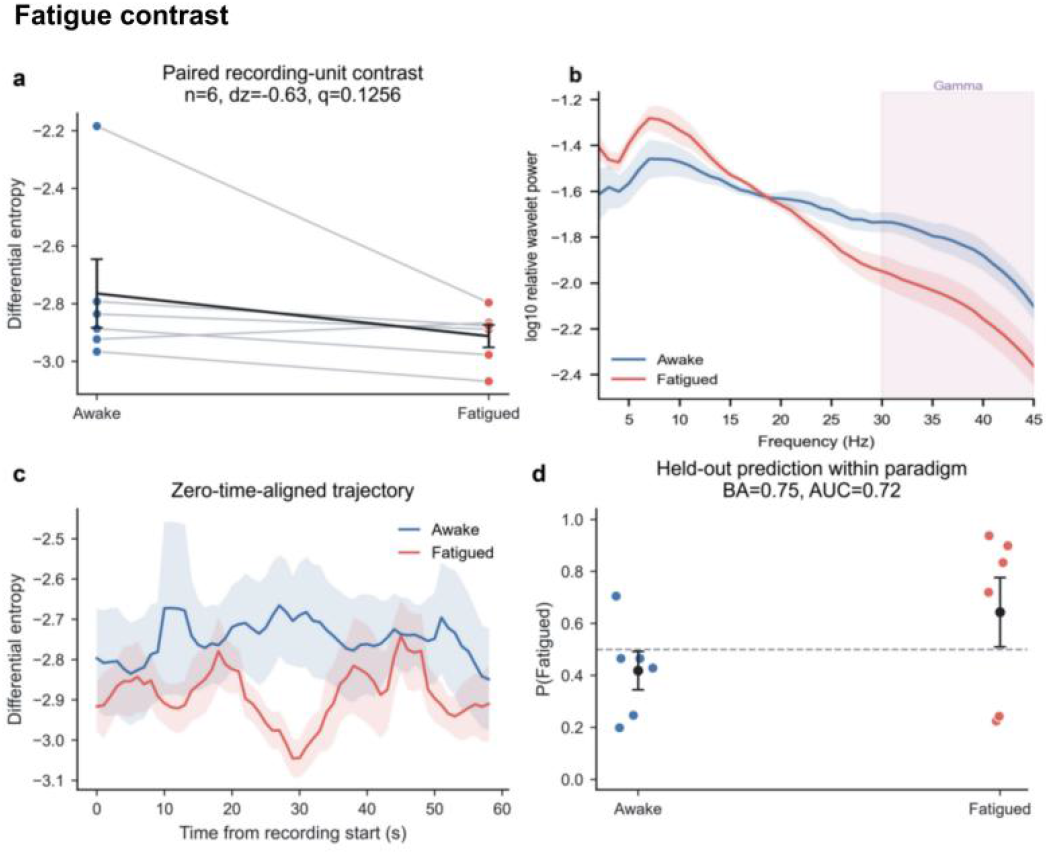
Fatigue contrast. (a) Paired differential entropy (n = 6; fatigued – awake mean difference, −0.1478; 95% CI, −0.3462 to −0.0158; dz = −0.63; sign-flip p = 0.1077; FDR q = 0.1256). (b) Within-paradigm power spectra, recording-unit mean log10 relative wavelet-power profiles for fatigue (n = 6);

### D. Attention Engagement

As illustrated in Fig. 7a, differential entropy exhibited elevated levels during periods of focused attention in comparison to those of the distracted state [27], a finding consistent across all six paired recording units. The mean focused-minus-distracted difference was 0.4013 (Table S1, 95% CI, 0.2629 to 0.5408), corresponding to a large standardized effect (Fig. 7a, dz = 2.02; Table S1, sign-flip P = 0.0462). Although the FDR-adjusted value exceeded the confirmatory threshold (Fig. 7a, q = 0.1077), the 6/6 directional agreement and positive confidence interval showed consistent condition sensitivity of the eyewear-based EEG system to detect attention engagement.

**FIG. 7.**
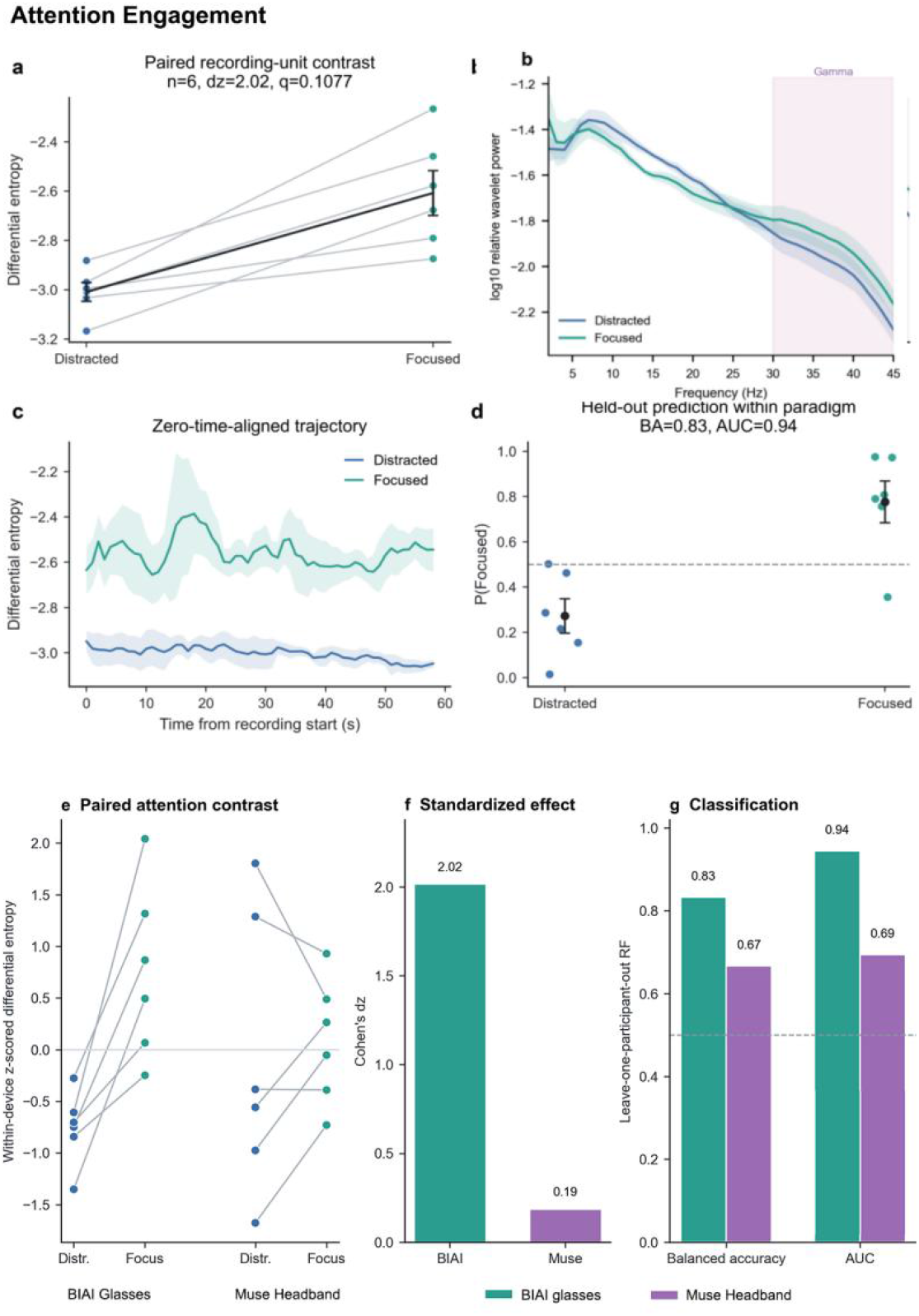
Attention engagement. (a) Paired differential entropy (n = 6; focused – distracted mean difference, 0.4013; 95% CI, 0.2629 to 0.5408; dz = 2.02; sign-flip p = 0.0462; FDR q = 0.1077). (b) Within-paradigm power spectra, recording-unit mean log10 relative wavelet-power profiles for attention (n = 6); ribbons show s.e.m. Gamma shading spans 30-45 Hz. (c) Zero-time-aligned trajectories. (d) Group-held-out predictions (balanced accuracy, 0.833; AUC, 0.944). Attention benchmark against Muse headband: (e) Within-device standardized differential entropy for distracted and focused recordings. (f) Standardized focused-minus-distracted effects (BIAI glasses of EEG device, dz = 2.02; Muse headband: dz = 0.19). (g) Leave-one-subject-out random-forest performance (BIAI EEG glasses: balanced accuracy, 0.833; AUC, 0.944; Muse headband: balanced accuracy, 0.667; AUC, 0.694). The devices were recorded in separate cohorts; raw amplitude-dependent entropy values were not compared directly.

At the single-band level, the focused-minus-distracted gamma effect was dz = 0.40 (Fig. 9), and no band met the corrected threshold (Figs. 7b and 9). However, condition differentiation was more evident in differential entropy (dz = 2.02, Fig. 7a and Table S1) than in any single wavelet band (Fig. 9). In addition, there was a sharply separated between distracted and focused attention in the zero-time-aligned trajectory (Fig. 7c), and significant multivariate prediction (balanced accuracy, 0.833; AUC, 0.944) (Fig. 7d, Table 1). Compared with that of the Muse headband [13], [44], the comprehensive directional agreement, the substantial standardized effect, and the held-out prediction were taken together to suggest a heightened sensitivity to attention manipulation in the recordings of our eyewear system (Fig. 7e-g). Moreover, it is further supported by a BCI function that our EEG glasses directly controlled the drone to take off, through the EEG signals captured from the user’s focused attention state (shown in Video S1).

### E. Emotion Recognition

A statistically significant difference in the levels of differential entropy was observed among negative, neutral, and positive recordings for emotion recognition (Friedman chi-squared = 23.41; p = 8.2 × 10^−6^; FDR q = 1.9 × 10^−5^) (Fig. 8a) [45]. The positive-minus-negative contrast was found to be positive in all 17 paired units (mean difference, 0.6787; 95% CI, 0.4149 to 1.0078; dz = 1.05; sign-flip p = 5.0 × 10^−5^; FDR q = 1.2 × 10^−4^) (Fig. 8b). In terms of zero-time-aligned trajectory (Fig. 8c), there was a dramatically separation between positive and negative emotion states, while the neutral one was separately located between them and a little closer to the negative one. Group-held-out classification distinguished positive from negative emotion with a balanced accuracy of 0.882, an F1 score of 0.889, and an AUC of 0.976 (Table 1, Emotion, 2 classes). Then, the three-class negative-neutral-positive model achieved a balanced accuracy of 0.667, a macro-F1 score of 0.669, and a one-versus-one macro AUC of 0.815 (Table 1, Emotion, 3 classes; and Fig. 8d).

**FIG. 8.**
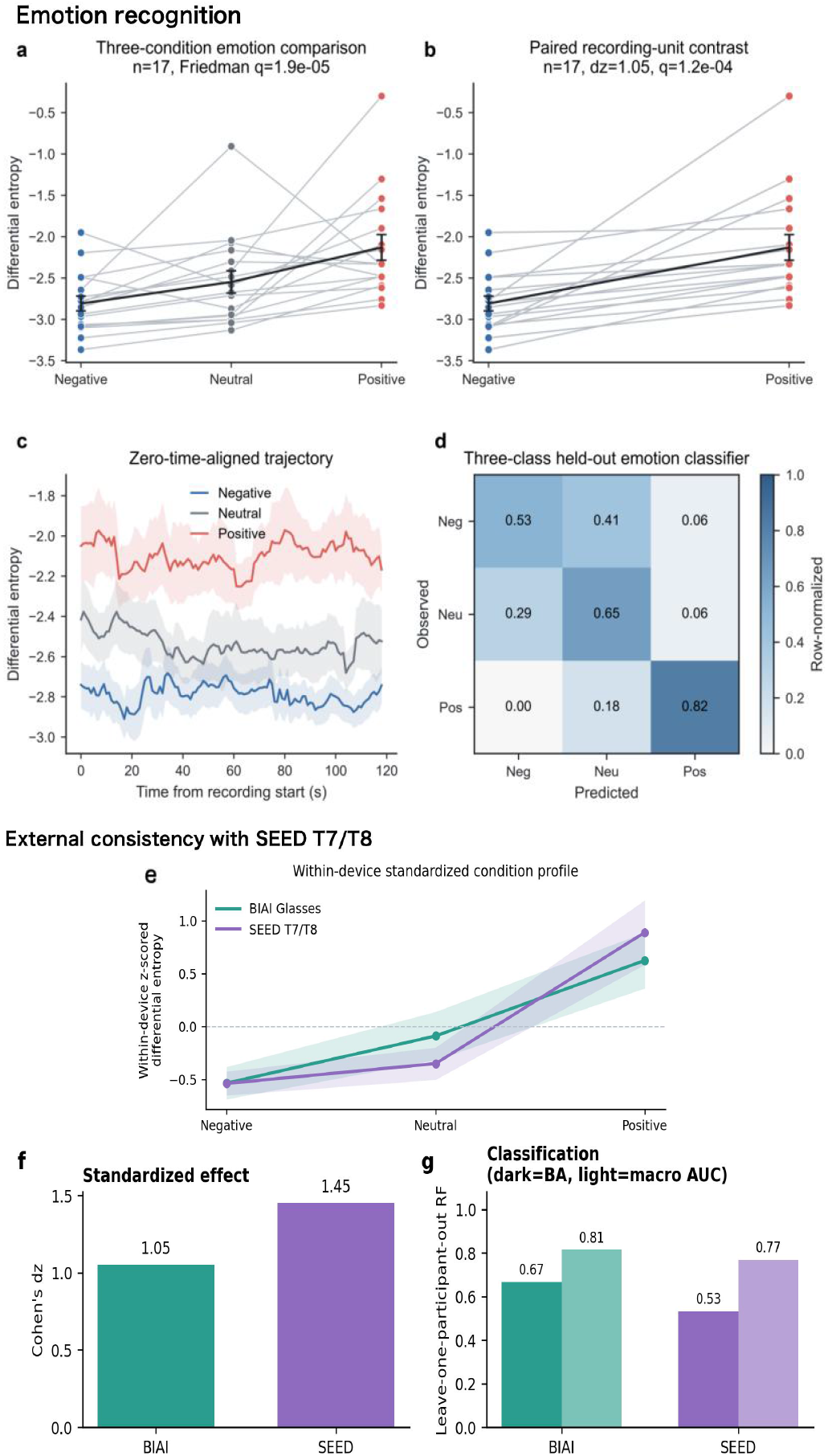
Emotion recognition. (a) Three-condition differential entropy (n = 17; Friedman chi-squared = 23.41; p = 8.2×10^−6^; FDR q = 1.9 ×10^−5^). (b) Positive-minus-negative contrast (mean difference, 0.6787; 95% CI, 0.4149 to 1.0078; dz = 1.05; sign-flip p = 5.0 × 10^−5^; FDR q = 1.2 × 10^−4^). (c) Zero-time-aligned trajectories. (d) Three-class held-out confusion matrix (balanced accuracy, 0.667; macro AUC, 0.815). (e) Within-device standardized differential entropy across negative, neutral, and positive conditions for BIAI Glasses and SEED T7/T8; values are z-scored within each dataset, so absolute DE values are not comparable. (f) Standardized positive-minus-negative effects (Cohen’s dz) for BIAI Glasses (1.05) and SEED T7/T8 (1.45). (g) Leave-one-participant-out random-forest classification for the three-class problem; dark bars show balanced accuracy (BA) and light bars show macro AUC (BIAI Glasses: BA = 0.67, macro AUC = 0.81; SEED T7/T8: BA = 0.53, macro AUC = 0.77).

**FIG. 9.**
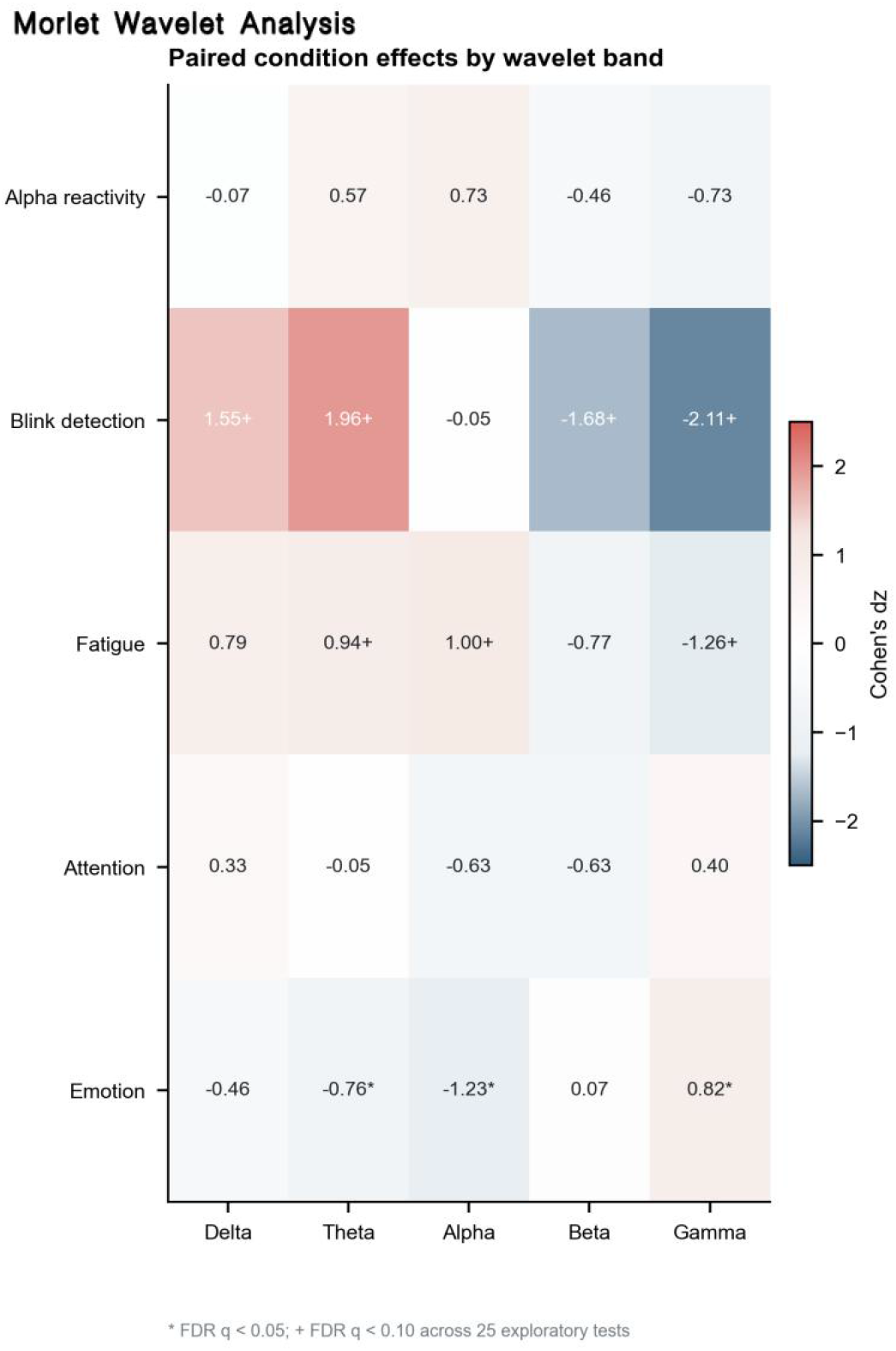
Morlet wavelet analysis. Paired Cohen ‘ s dz across five tasks and five frequency bands. Asterisks (*) indicate FDR q < 0.05 and plus signs (+) indicate q < 0.10 across 25 exploratory tests.

To test spatial consistency with a conventional temporal-lobe reference, we re-analyzed the SEED dataset using only T7 and T8 sites [28]. The positive-minus-negative differential entropy contrast was positive in all 15 participants (mean difference = 0.4237; 95% CI [0.2921, 0.5762]; dz = 1.45; sign-flip p=1.0 ×10^−4^). T7 and T8 analyzed separately showed similar effects (dz = 1.35 and 1.31). Under leave-one-participant-out random-forest validation, SEED T7/T8 achieved a balanced accuracy of 0.767 and an AUC of 0.924 for positive-versus-negative classification, and 0.533 and 0.769 for three-class classification. Then our eyewear system showed the same positive > negative direction across 17 recording units (dz=1.05; 17/17 positive; FDR q=1.2×10^−4^). These findings further support a T7/T8-consistent valence direction, might as the best sites to detect emotional valence at the electrophysiological level (Fig. 8e-g, Table S2), regardless of portable or conventional EEG devices [28], [34], [53].

Therefore, the consistent increase in differential entropy from negative to positive clips, the corrected three-condition effect, and the held-out prediction converged on systematic differentiation of video-elicited emotion states. Among the available evidence of our eyewear system (Table 1, S1 and Fig. 9), moreover, that pertaining to emotion was the strongest in terms of its state-related characteristics. As shown in Fig 9, there was a significantly increased gamma-band activity reflected on positive valence [50], also with decreased activities of alpha and theta bands in the morlet wavelet analysis (FDR * q < 0.05, dz = 0.82, for gamma band; dz = −1.23 and −0.76, for alpha and theta band) (Fig. 9a).

## V. Discussion

This present study evaluates the eyewear-based EEG system across five EEG validation paradigms: alpha oscillation with closed eyes, voluntary blinking, fatigue-related resting state, focused attention, and video-elicited emotion. The resulting data provides feasibility evidence for eyewear-based EEG system acquisition. The primary contributions of this study are enumerated below: At the technical level, we address part of the portability and setup burden associated with traditional EEG devices. We propose a lightweight EEG acquisition solution based on dry electrodes made of titanium materials, inferoposterior to the T7/T8 sites without fair covering around the ears. This solution is integrated into a glasses frame, as shown in Fig. 1. This design with glasses frame enhances structural integration and user experience [54], [55], potentially serving as a model for future miniaturized wearable EEG systems [56]. At the level of application, the device was evaluated across series of common scenarios pertaining to the utilization of EEG. Through the execution of experiments involving multiple tasks, we systematically explored the applicability of the device across various fields of EEG.

Moreover, it was demonstrated that our eyewear system can control the drone to roll and take off directly through the EEG signals with blinking and focused action (Video S1 and S2), respectively. This exploration supports the potential of the eyewear system to complement conventional EEG in selected low-burden monitoring scenarios.This not only offers reliable hardware support for bringing EEG technology from the laboratory into daily life, but also provides a novel technical path for mental health management with EEG detection of fatigue (Fig. 6), attention (Fig. 7), and emotion (Fig. 8) via eyewear system. It is worthy of providing more data support for its potential in home health monitoring and daily medical application in near future, [17], [57].

The present results indicate that our EEG glasses retained usable recordings across paradigms and showed the clearest evidence for blink sensitivity (Fig. 5) and emotion recognition (Fig. 9), whereas alpha reactivity, fatigue, and attention remain exploratory in the current samples (Table 1 and S1). These findings may be indicative of fundamental hardware design and signal acquisition mechanisms of this eyewear-based EEG system [18], [57]. Thus, the ensuing discourse will delve into the three pivotal design aspects, including electrode site selection, reference electrode layout, and frequency band adaptability. This will serve to elucidate the underlying rationale for the observed signal patterns and to delineate the boundaries of the current validation [58].

The temporal-lobe-adjacent electrode placement facilitates the capture of signals relevant to multiple task states. One plausible explanation for our data is the functional relevance of temporal regions and the specificity of certain EEG signals to the tasks evaluated in this study. From an anatomical and neurophysiological perspective, the T7 and T8 sites correspond to the part of the temporal lobes of the brain [24], [59], while the EEG signals captured by the subject’s glasses temples are closer to the inferior temporal gyrus (Fig. 1). The temporal lobes exhibit extensive neural connections with other cortical regions, including the frontal and occipital lobes. This region is among the key center for cognitive functions involving multisensory integration [59], attention processing [14], [60], and emotion encoding with memory [61]. These functions are pertinent to several paradigms in this study, particularly affective-state differentiation or emotion recognition (Fig. 8 and 9). However, the evidence present design does not further isolate anatomical source yet. This placement is most likely, therefore, contributing to the observed within-paradigm feature differences. According to the findings of preceding studies, T7/T8 signals recorded by some portable EEG devices were comparable to temporal electrode signals recorded with traditional multi-channel EEG caps [26], [27], [28], effectively preserving the core features in terms of amplitude and frequency [28], [53], [61],. Furthermore, those EEG signals involved T7/T8 sites have provided electrophysiologically key values associated with some common brain disorders, including major depressive disorder [62], [63], Alzheimer’s disease [64], attention-deficit/hyperactivity disorder [65], and temporal lobe epilepsy as well [66]. The T7/T8-consistent direction observed in the SEED re-analysis is physiologically plausible given the proximity of the temple contacts to temporal lobes involved in affective processing (Fig. 8 e-g, and Table S2) [28], [34]. Future work will further record the same participants with an eyewear-based EEG system and a multichannel cap system in a synchronized or crossover design.

Our EEG glasses frame utilizes the nasal bridge as the reference electrode (Fig. 1a), offering a compact and user-friendly reference configuration. The fundamental principle underlying EEG acquisition is the measurement of electrical potential differences between two points on the scalp. The reference electrode has to provide a stable potential baseline to minimize interference. Its placement was to directly impact signal stability and cost-effectiveness, as well as the extraction of EEG features in various scenarios [31]. Then, the nasal bridge area exhibits inherent advantages [30], [31]: firstly, the skin is thin, with minimal fat and low impedance, enabling effective electrical contact with the reference electrode. The implementation of stable reference measurements has been demonstrated to be achieved with the nose-pads made of titanium materials (Fig. 4-8), thereby reducing the complexity and cost associated with equipment usage (Fig. 1). Secondly, the location of the nasal bridge at the center of the face, a region that is distant from the primary brain activity regions, results in minimal physiological electrical activity. This configuration may offer a relatively stable reference location and reduce setup burden; however, signal-to-noise comparisons are still needed to further quantify its electrophysiological advantage. While compared with conventional reference electrodes, such as earlobes and mastoids, the bridge of the nose is situated at a further distance from the T7 and T8 recording sites, and this configuration enables the precise capture of potential differences. Concurrently, the integration of the reference electrode into the nose bridge of the glasses frame serves to streamline the structural design, thereby achieving a balance between the device’s lightweight construction and its recording performance.

The spectral panels suggest that higher-frequency activity may contribute to separation in several paradigms, especially the gamma band (30-45 Hz) for recognizing states of fatigue (Fig. 6b), attention (Fig. 7b), and emotion (Fig. 9). From a physiological perspective, the gamma frequency band is primarily linked to higher cognitive functions of the brain [34]. and its activity intensity directly corresponds to the level of synchronization of neuronal populations involved in cognitive and emotional processes [24], [50]. Furthermore, given the established role of the T7 and T8 sites in temporal lobes as critical regions for cognitive functions, their heightened gamma-band activity is particularly evident [50], [67]. As shown in Fig. 6b, this study reveals that fatigue results in a substantially more pronounced decline in the gamma band compared to other bands. With regard to the detection of attention, as illustrated in Fig. 7b, the integration of target information requires the high synchronization of neuronal populations [60], accompanied by a substantial increase in gamma-band activity detected by our eyewear system. In the context of emotion recognition, widespread cortical activation induced by positive emotions has been shown to promote a substantial increase in gamma-band activity. The increased gamma-band activity can serve as an accurate reflection of positively emotional changes, making it the most discriminative frequency band (dz = 0.82), with lower activities of alpha and theta bands (FDR * q < 0.05, Fig. 9). Nevertheless, future study should test band-specific contributions using pre-specified features, larger paired samples, and behavioral labels.

In general, our eyewear-based EEG system provides a wearable, low-burden route for recording EEG signals from both sites inferoposterior to T7/T8 with a metal nose-pad reference. The extant data supports the feasibility of our eyewear system as an EEG acquisition solution, as well as its capacity for high epoch retention, great sensitivity to voluntary blinking, and controlled emotion recognition. The evidence for fatigue and attention monitoring remains exploratory, and direct comparisons with conventional EEG caps and long-duration real-world recordings are still required. Future research would aim to augment the sample sizes, incorporate more behavioral labels with clinical categories.

## VI. Conclusion

In this study, we proposed a solution of eyewear-based EEG system and evaluated its reliability and validity for EEG acquisition. The objective of this study was to extend the applications of EEG beyond laboratory conditions. The EEG glasses frame exhibited a wearable and user-friendly design, featuring a classical frame constructed from nylon and titanium, conventional materials in the fabrication of eyewear. Subsequent experiments yielded evidence substantiating the superior quality of EEG signals captured by our advanced eyewear system. These findings suggest that eyewear-based EEG system with metal temples and nose pads can reliably capture EEG signals across a range of physiological and cognitive states. Therefore, this device may be considered appropriate for daily use in the real-world, such as the management of mental health, with the EEG signals captured around the ears inferoposterior to T7/T8 sites. In the future, this eyewear-based EEG system could be more widely evaluated by diverse participants, including those with brain disorders, and their algorithms could be further optimized with additional data.

## VII. Acknowledgment

We would like to express our profound gratitude to all participants for their invaluable contributions to this research.

