## Supplemental materials for "An Eyewear-Based EEG Interface for Temporal-Region Monitoring and Emotion Recognition"

**Legends and Figures**

Video S1

Video S2

Figure S1

Table S1

Table S2

Video S1. A BCI function controls the drone to take off, with the EEG signals of attention state, captured by our EEG smart glasses.

1.mp4

Video S2. A BCI function to control the roll action of drone ball is caused by blinking EEG signals using our EEG smart glasses.

2.mp4

Figure S1

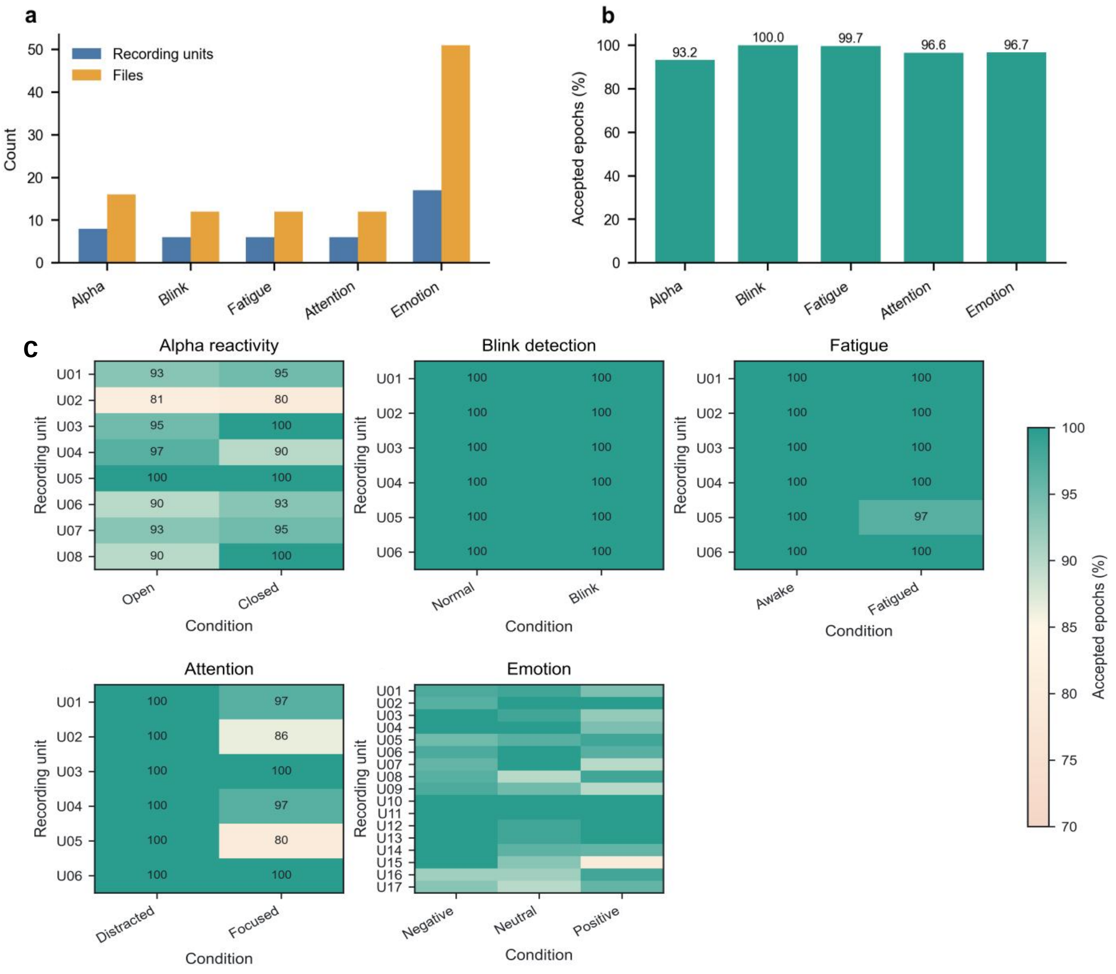


**FIGURE S1.** Dataset structure and signal quality. (a) Recording-unit and file counts after addition of four eyes-open/eyes-closed pairs. (b) Mean retained epochs after within-recording artifact control. (c) Recording-level epoch acceptance. Rows denote recordings and columns denote sequential 2-s epochs. Colour indicates retained and rejected epochs after within-recording quality control. the epoch-retention matrix was retained as an audit trail for preprocessing. It shows where each recording contributed accepted or rejected epochs after robust quality control. This information is useful because dry-electrode wearable EEG can be affected by transient contact changes, and transparent retention reporting helps distinguish physiological conclusions from preprocessing artifacts.

Table S1

### TABLE S1

PARADIGM-SPECIFIC PRIMARY CONTRASTS. P VALUES ARE TWO-SIDED SIGN-FLIP TESTS; Q VALUES ARE BENJAMINI-HOCHBERG ADJUSTED WITHIN EACH PARADIGM-SPECIFIC FEATURE FAMILY.

| Paradigm | Contrast | Primary feature | n | Mean difference | 95% CI | dz | P | q |
|---|---|---|---|---|---|---|---|---|
| Alpha oscillation | Closed - open | Alpha/beta ratio | 8 | 0.3893 | 0.0962 to 0.7152 | 0.80 | 0.0661 | 0.2588 |
| Blinking detection | Blink - normal | Peak-to-peak | 6 | 0.3404 | 0.1983 to 0.4741 | 1.76 | 0.0462 | 0.0497 |
| Fatigue contrast | Fatigued - awake | Differential entropy | 6 | -0.1478 | -0.3462 to -0.0158 | -0.63 | 0.1077 | 0.1256 |
| Attention engagement | Focused - distracted | Differential entropy | 6 | 0.4013 | 0.2629 to 0.5408 | 2.02 | 0.0462 | 0.1077 |
| Emotion recognition | Positive - negative | Differential entropy | 17 | 0.6787 | 0.4149 to 1.0078 | 1.05 | 5.0e-05 | 1.2e-04 |

Table S2

Table S2

External consistency between the EEG smart glasses and SEED T7/T8 re-analysis.

| Measure | Wearable EEG glasses | SEED T7/T8 |
|---|---|---|
| Task | Video-evoked three-class emotion | Video-evoked three-class emotion |
| Primary contrast | Positive minus negative DE | Positive minus negative DE |
| Sample | 17 recording units | 15 participants |
| Mean difference | 0.6787 device units | 0.4237 SEED internal units |
| 95% CI | — | [0.2921, 0.5762] |
| Cohen’s dz | 1.05 | 1.45 |
| Directional agreement | 17/17 positive | 15/15 positive |
| Sign-flip p | 5.0×10−55.0×10−5 | 1.0×10−41.0×10−4 |
| FDR q | 1.2×10−41.2×10−4 | — |
| Positive vs negative: BA/AUC/F1 | 0.882 / 0.976 / 0.889 | 0.767 / 0.924 / 0.759 |
| Three-class: BA/macro AUC/macro F1 | 0.667 / 0.815 / 0.669 | 0.533 / 0.769 / 0.520 |

Note: Raw DE values and absolute amplitudes are not comparable across devices. Comparison is limited to direction, within-device standardized effects, and participant-level held-out metrics.
